# Development, validation and application of an LC-MS/MS method quantifying free forms of the micronutrients queuine and queuosine in human plasma using a surrogate matrix approach

**DOI:** 10.1101/2024.06.24.600019

**Authors:** Xiaobei Pan, Swathine Chandrasekaran, Jayne V. Woodside, Steffi G. Riedel-Heller, Martin Scherer, Michael Wagner, Alfredo Ramirez, Brian D. Green

## Abstract

Queuosine (Q) is a hypermodified 7-deaza-guanosine nucleoside exclusively synthesized by bacteria. This micronutrient and its respective nucleobase form queuine (q) are salvaged by humans either from gut microflora, or digested food. Depletion of Q-tRNA in human or mouse cells causes protein misfolding that triggers endoplasmic reticular stress and activation of the unfolded protein responses. In vivo, this reduces neuronal architecture of the mouse brain affecting learning and memory. Herein, a sensitive method for quantifying free q and Q in human blood was developed, optimised and validated.

After evaluating q/Q extraction efficiency in several different solid-phase sorbents, Bond Elut PBA (phenylboronic acid) cartridges were found to have the highest extraction recovery for q (82%) and Q (71%) from pooled human plasma. PBS with 4% BSA was used as surrogate matrix for method development and validation. An LC-MS/MS method was validated across the concentration range of 0.0003 – 1 µM for both q and Q, showing excellent: linearity (r^2^ = 0.997 (q) and r^2^ = 0.998 (Q)), limit of quantification (0.0003 µM), accuracy (100.39% - 125.71%) and precision (CV% < 15.68%). In a sampling of healthy volunteers (n = 44) there was no significant difference in q levels between male (n = 14; mean = 0.0068 µM) and female (n = 30; mean = 0.0080 µM) participants (p = 0.50). Q was not detected in human plasma. This validated method can now be used to further our understanding of the role of q/Q in nutrition, physiology and pathology.

## Introduction

It has been reported that over 170 RNA modifications in RNA molecules occur across all domains of life (1). Amongst all the RNA molecules, tRNA is the most extensively modified RNA, containing on an average 13 modifications per molecule (2). tRNA modifications can direct mRNA decoding when it occurs in the anticodon loop (position 34). Modifications occurring downstream of the anticodon nucleotides (position 37) affect the stability, folding, localization and translational dynamics of tRNA (2, 3). Among all the tRNA modifications, queuosine (Q) is the only nucleoside that is synthesized in bacteria *de novo* and salvaged in eukaryotes from their environment (4). It is a hypermodified 7-deaza-guanosine nucleoside that replaces the unmodified guanine at wobble position 34 of the anticodon GUN for four tRNAs acceptors for the amino acids tyrosine, asparagine, aspartic acid and histidine (5, 6). The Q biosynthesis pathway has been well studied in eubacteria (7, 8). However, in eukaryotes the biosynthesis pathways and cellular roles of Q remain to be fully elucidated. Humans obtain Q as a micronutrient from diet or gut microflora. An enzyme called QNG1 (previously known as DUF2914) hydrolyses Q or a Q monophosphate to release queuine (q), which in turn can be salvaged and reincorporated into tRNA by the eukaryotic tRNA guanine transglycosylase (eTGT) which is comprised of the QTRT1 and QTRT2 subunits (9).

It has been demonstrated that the depletion of Q-tRNA in both mouse and human cells leads to protein misfolding, resulting in endoplasmic reticular stress and activation of the unfolded protein response (10, 11). The levels of tetrahydrobiopterin (BH4) are decreased when human HepG2 cells are deprived of q, and also in mice with an engineered deficiency of Q-tRNA (through the disruption of the TGT enzyme) (12). BH4 is a critical cofactor in the production of numerous biogenic amine neurotransmitters, and this has led to speculation that Q-tRNA status could be an important factor in neurological and neuropsychiatric diseases (13). Indeed, the administration of an analog of q led to full remission of multiple sclerosis, by reducing effector immune cell proliferation and activation, in a mouse model of this disease (14). Furthermore, the addition of synthesised q to an in vitro model of Alzheimer’s disease and Parkinson’s disease indicates that it has neuroprotective effects (15). Recently, Q-tRNA modification was found to occur ubiquitously across all mouse tissues, but tissues with the highest levels of modification where heart, brain and skeletal muscle (16). Perhaps the most convincing evidence of Q’s physiological importance so far, has been observed in the brain. The disruption of the TGT enzyme complex in mice to, prevent Q-tRNA modification, leads to alterations in the structure of hippocampal pyramidal neurons, reduction in cell density and global imbalance in the speed of codon-biased translation. This appears to impact on learning and memory formation in the mouse, which is particularly pronounced in females (16).

The two commonly used techniques to measure Q-tRNA modification are, firstly, acryloylaminophenyl boronic acid (APB) gel electrophoresis and secondly, LC-MS/MS analysis. In the presence of APB, the cis-diol group on Q slows the migration of Q-tRNA, therefore separating the Q-tRNA from other unmodified tRNA (17). Then the Q-tRNA band can be quantified by northern blot (18). The APB gel technique can directly detect Q-tRNA in individual tRNAs but obviously does not detect free q or Q which are more likely to circulate in blood. LC-MS/MS is another sensitive and accurate technique for measuring the molecular modification a large variety of biological materials including RNA (19-21). First, the RNA is isolated and purified from biological samples. Then, the nucleosides and nucleotides are enzymatically hydrolysed from the RNA, and individually quantified using LC-MS/MS analysis (22). Although these methods have previously been applied for Q-tRNA modification analysis, no validated LC-MS/MS method for extracting and quantifying free q and Q in complex biological samples has yet been reported.

The aim of this study was therefore to establish an accurate and reliable LC-MS/MS method for quantifying free q and Q in blood samples and to assess the influence of blood collection and processing conditions in its measurement.

## Methods

### Materials

Q standard was provided by Professor Vincent Kelly (Trinity College Dublin, Ireland) and a Q stock solution (1mM in water) was provided by Professor Valerie de Crecy-Lagard (University of Florida, USA). Indomethacin, butylated hydroxytoluene (BHT), diethylenetriamine pentaacetate (DTPA), sodium carbonate, sodium hydroxide, phosphate buffered saline (PBS), ammonia, ammonium acetate and methanol were purchased from Merck (Missouri, USA). Serum, EDTA and lithium heparin blood collection tubes were purchased from BD Vacutainers. LC-MS grade acetonitrile, methanol and formic acid (FA) were purchased from Honeywell (North Carolina, USA).

### Subjects

To optimise the experimental protocol for blood collection, processing and extraction, blood samples were obtained from volunteers at the Centre for Public Health, Queen’s University Belfast. Ethical approval was not required for these method development studies and the blood samples were anonymised and pooled.

To assess q/Q levels across a range of healthy participants, plasma samples (n = 44) were accessed from the biobank of the German study on Aging, Cognition and Dementia (AgeCoDe) (23). The study ethical approval and study protocol are detailed in the Supplementary material.

### Preparation of stabilization cocktail

A stabilization cocktail was prepared using indomethacin (20 mM in 5%NaHCO_3_ and then diluted to 0.2 mM with 1xPBS), BHT (5 mM in ethanol), and DTPA (10 mM in PBS, pH 7.4 adjusting with 6N NaOH firstly and then with 1N NaOH). The cocktail was added to the collection tubes before blood collection composed of per 1 mL of blood: 75 µL of indomethacin, 4 µL of BHT and 10 µL of DTPA.

### Blood samples collection

For blood collection method optimization, blood samples from 3 volunteer participants were collected using EDTA, lithium heparin and serum tubes. One set of samples was treated with stabilisation cocktail, kept at 4°C for 30 min, and then centrifuged at 2400 rpm for 20 minutes at 4°C. Untreated EDTA and lithium heparin samples were centrifuged immediately after collection. Untreated serum samples were allowed to clot before centrifugation. The stabilization cocktail was prepared freshly on each day of collection. The three blood samples collected in the same tube were then pooled before q/Q extraction.

### Selection of SPE cartridges and sample purification

Six types of SPE cartridges, including Oasis HLB, Oasis MCX, Oasis MAX and Oasis SAX, EBVI CARB and Bond Elut PBA were selected for purification treatment. Oasis HLB, Oasis MCX, Oasis MAX and Oasis SAX were obtained from Waters (Milford, USA). ENVI CARB was purchased from Merck (Darmstadt, Germany). Bond Elut PBA (phenylboronic acid) cartridge and 96-well plate were purchased from Agilent (Santa Clara, USA). 1 µM of q and Q was spiked in human plasma and extracted with SPE cartridges. The detailed extraction procedures can be found in Supplementary Table 1.

### Preparation of calibration standards and quality controls

Queuine standard was dissolved in ultrapure water to prepare 1 mM stock solution. Phosphate buffered saline (PBS) with 4% bovine serum albumin (BSA) was sued as surrogate matrix. Both q and Q stock solutions were aliquoted and stored at -20 ?C. Calibration standards (1, 0.3, 0.1, 0.03, 0.01, 0.003, 0.001 and 0.0003 µM), and 3 quality controls (QCs; 0.8, 0.02 and 0.0008 µM) were prepared freshly either in ultrapure water; or spiked in matrix surrogate and then extracted using a PBA cartridge on the day of analysis.

### Extraction of q and Q

For spiked samples, 500 µL of surrogate matrix was mixed with 50 µL of spiking standards and 450 µL of ammonia solution (1M, pH 11.35). In case of study samples, 500 µL of blood mixed with 500 µL of ammonia solution (1M, pH 11.35). The Bond Elut PBA 1cc column was conditioned with 1 mL of acetonitrile-water (3:7, v/v) with 1% formic acid and equilibrated with 1 mL of ammonia solution (1M, pH 11.35). The diluted samples were loaded onto the column, followed by washing with 1 mL of 5% acetonitrile at pH 11.35. The analytes were eluted using acetonitrile-water (3:7,v/v) with 5% methanol and 1% formic acid. The elute was then vacuum dried and reconstituted in 100 µL of water with 5% methanol. Similarly, for cohort samples, a 1ml PBA 96-well plate was used, and the volume of serum samples used was reduced to 200 µL and diluted using 200 µL of ammonia solution (1M, pH 11.35) before processing through the plate.

### Chromatographic and mass spectrometric conditions

LC-MS/MS analysis was performed on a system consisting of an AB SCIEX ExionLC system (Foster, USA) coupled to an AB SCIEX Triple Quad 5500+ mass spectrometer (Foster, USA). The mass spectrometer was operated in ESI positive mode using Multiple Reaction Monitoring (MRM). Chromatographic separation was achieved with a Waters XSelect HSS T3 column (100 × 4.6 mm, 3.5 μm, Milford, USA) at 45°C using a gradient of 0.1% formic acid in water (mobile phase A) and 0.1% formic acid in acetonitrile (mobile phase B) at a flow rate of 1.0 mL/min. The mobile gradient and mass spectrometric parameters were set as previously described (24).

### Method validation

The established UPLC-MS/MS method was evaluated and validated for linearity, sensitivity, matrix effect, recovery, and precision. Recovery and precision were evaluated by spiking q or Q into 4% BSA PBS at three different concentrations.

#### a) Linearity and sensitivity

Linearity was evaluated by the 8-points calibration curve of q and Q in the range of 0.3 nM – 1.0 µM. The sensitivity of method was assessed by the limit of detection (LOD) and limit of quantitation (LOQ) for q and Q at signal-to-noise (S/N) ratio of 3 and 10, respectively.

#### b) Accuracy and precision

Five replicated sets of QCs (0.0008, 0.02 and 0.8 µM) were prepared in a surrogate matrix to determine the accuracy and precision of the assay. These experiments were carried out intraday (same day) and inter-day (after 24 h) during 5 days. The accuracy was calculated by [(the measured concentration)/(the spiked concentration)] x 100%, while the precision of the method was expressed by the coefficient of variation [(standard deviation)/mean] x 100%.

#### c) Extraction recovery, matrix effects and parallelism

Extraction recovery was measured by comparing the response of analytes added to extracted surrogate matrix (post-extraction spiked samples) and responses of analytes added to surrogate matrix before extraction (pre-extraction spiked samples).

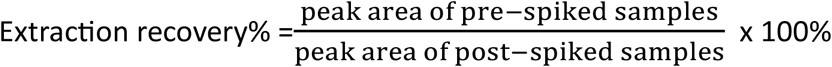

The assessment of on the ionization suppression/enhancement were carried out by comparing the response of analytes added to extracted surrogate matrix (post-extraction spiked samples) and the response of analytes in solvent.

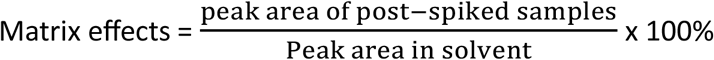

Evaluation of parallelism is of critical importance for endogenous analytes when surrogate matrix is applied for quantification. Batter of parallelism indicates that there is no significant difference between the surrogate and the authentic matrix. Two sets of curves were prepared either in surrogate matrix (4% BSA in PBS) or in human plasma with q and Q. Both sets of curves were prepared following the same preparation procedure. After LC-MS/MS analysis, linear regression equations were generated employing a linear regression model in each matrix, plotted using Microsoft Excel® and the relative percentage difference (RPD) were calculated.

#### d) Carryover

Carryover was evaluated by assaying solvent blank samples injected after the analysis of a surrogate matrix spiked with the highest QC.

### Application of the method to clinical human plasma samples

The validated method was applied to quantify q and Q in bio-banked human EDTA plasma from healthy volunteers. Mann-Whitney test was applied to determine if the levels of q and Q are significantly different between male and female participants.

## Results

### Selection of SPE cartridge

Six SPE cartridges, Oasis HLB, Oasis MCX, Oasis WCX and Oasis SAX, EBVI CARB and Bond Elut PBA, were selected to investigate the extract efficiency of spiked q and Q from human plasma (Fig. 1). The results showed that the Bond Elut PBA cartridge had the best extraction ability with the recovery of 82% for q and 71% for Q, followed by the Oasis MCX cartridge with the recovery of 44.5% for q and 54% for Q.

**Fig. 1.**
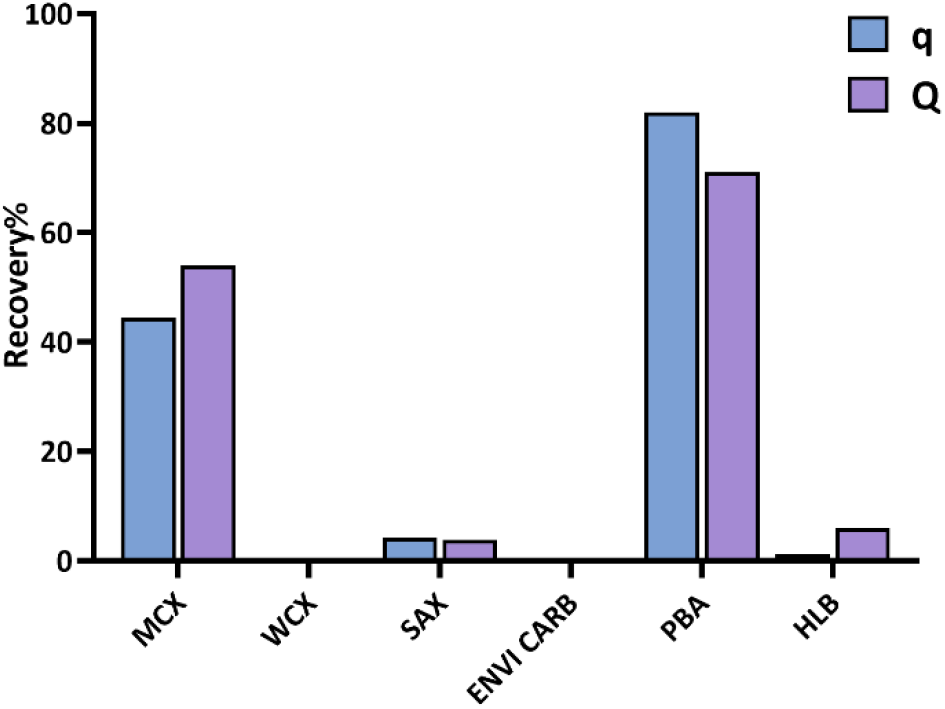
Recovery of q and Q extracted by different SPE cartridges

### Selection of blood collection tubes and inclusion of stabilization cocktail

Three blood collection tubes with or without stabilization cocktail were applied to evaluate the interference of anticoagulant and stabilization cocktail in the LC-MS/MS measurement. It was found that the EDTA plasma collected without stabilization cocktail obtained highest average q levels, while the serum with/without stabilization cocktail showed lowest experimental variability during the q/Q extraction procedure (Fig. 2). Adding stabilization cocktail did not improve the quantifiable levels of q. Thus it is unnecessary to add stabilization cocktail during blood collection. Serum samples are the first preference for measurement for q in future studies, and the next preference would be EDTA plasma without any stabilization cocktail.

**Fig. 2.**
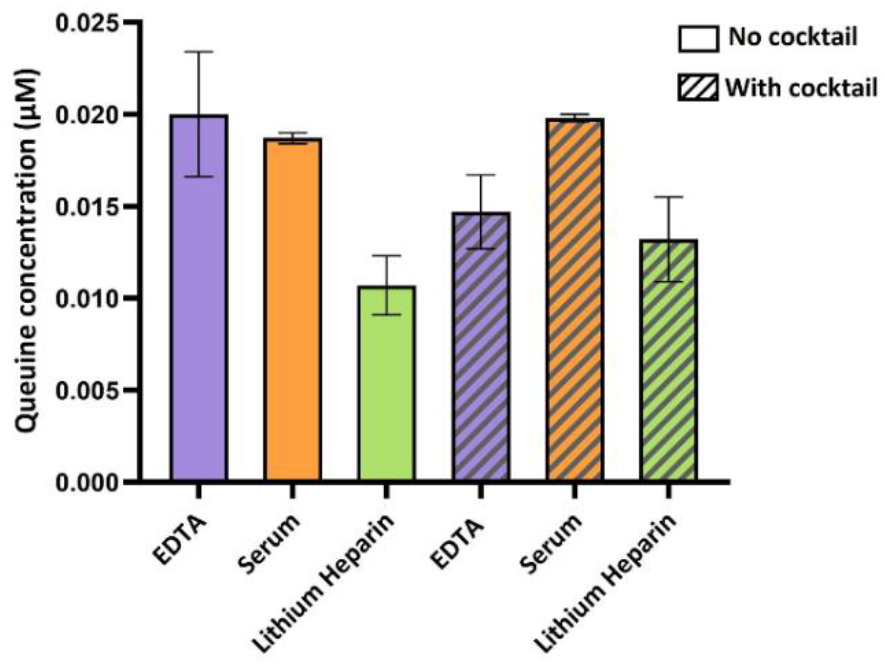
Measured concentration of q in human blood collected using different tubes with/without stabilization cocktail

### Validation of LC-MS/MS methods

#### a) Linearity and sensitivity

The calibration curves were generated at 8 concentration levels. It was shown that the linearity was good in the range from 0.3 nM to 1 µM. The mean regression coefficients (r^2^) for q is 0.997 ± 0.002 (n = 5) and for Q is 0.998 ± 0.003 (n = 5). The LOD and LOQ of the developed method were 0.1 nM and 0.3 nM, respectively, for both q and Q.

#### b) Accuracy and precision

The intra- and inter-assay accuracies and precisions for surrogate matrix are shown in Table 1. For q, the intra-day assay accuracy of the QCs in surrogate matrix ranged from 100.39% – 119.48% with %CV less than 12.24% across all levels. The inter-assay accuracy of QCs in surrogate matrix ranged from 104.28% – 113.41% with %CV less than 9.38% across all levels. For Q, the inter-assay accuracy of surrogate QCs ranged from 103.91% – 125.71% with %CV less than 15.68% across all levels. The inter-assay accuracy of surrogate QCs ranged from 104.89% – 124.67% with %CV less than 14.69%.

**Table 1.**
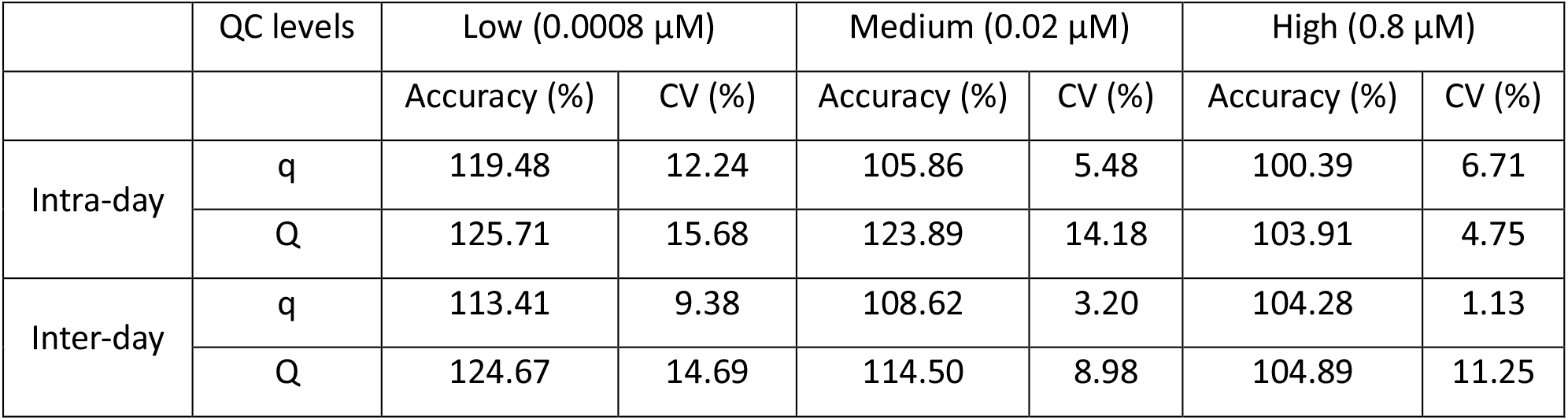
Intra-day and inter-day accuracy and precision of the QC samples.

#### c) Matrix effects, extraction recovery and parallelism in surrogate matrix

For both q and Q, the matrix effect at the low QCs exceeded 200%, which indicated the ionization enhancement by the coeluting undetected matrix components for the low QC (Table 2). The recovery was assessed to evaluate the extraction efficiency of analytes. For q, the extraction recoveries are all higher than 70%. However, for Q, the recoveries are lower than 50%, which indicates loss of Q in the solid phase extraction (Table 2).

**Table 2.** Matrix effects and extraction recovery of the QC samples.

| | Low (0.0008 $\mu\text{M}$) | | Medium (0.02 $\mu\text{M}$) | | High (0.8 $\mu\text{M}$) | |
| --- | --- | --- | --- | --- | --- | --- |
|  | Matrix effect (%) | Recovery (%) | Matrix effect (%) | Recovery (%) | Matrix effect (%) | Recovery (%) |
| q | 242.06 ± 5.23 | 71.43 ± 5.90 | 137.97 ± 3.70 | 76.20 ± 8.13 | 105.40 ± 1.59 | 74.87 ± 3.07 |
| Q | 280.23 ± 14.33 | 47.35 ± 15.88 | 171.87 ± 3.96 | 37.92 ± 6.20 | 128.93 ± 2.77 | 40.61 ± 2.82 |

Parallelism was performed to evaluate the equivalence of the surrogate matrix (4%BSA in PBS) to the authentic matrix (human plasma) for q and Q (Figure 3). Both sets of curves were prepared following the same preparation procedure. The RPD of the slopes between human plasma and surrogate matrix curves for q and Q was 6.61% and 4.89% respectively. Currently there is no regulatory guidance on acceptance criteria for parallelism between the authentic and matrix calibration curves, most published guidance suggests that if the slopes from the two sets of curves with the RPD below 15%, the curves are considered to be adequately parallel (25, 26).

**Fig. 3.**
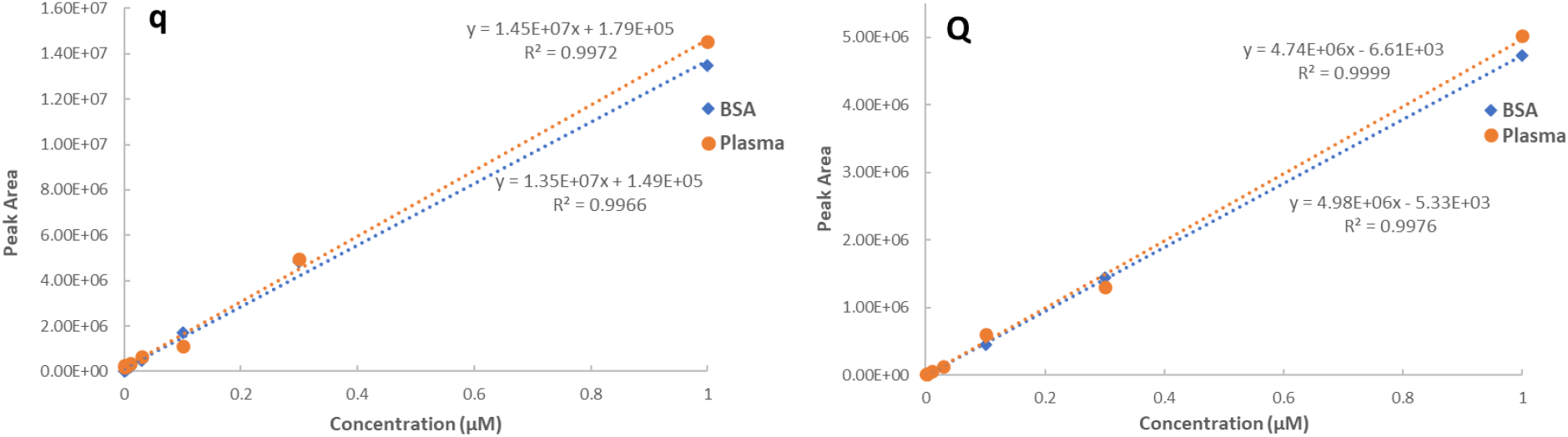
Parallelism of q (left) and Q (right) quantitation in surrogate matrix (4% BSA in PBS) and authentic matrix (human plasma) at their respective curve ranges

**Fig. 4.**
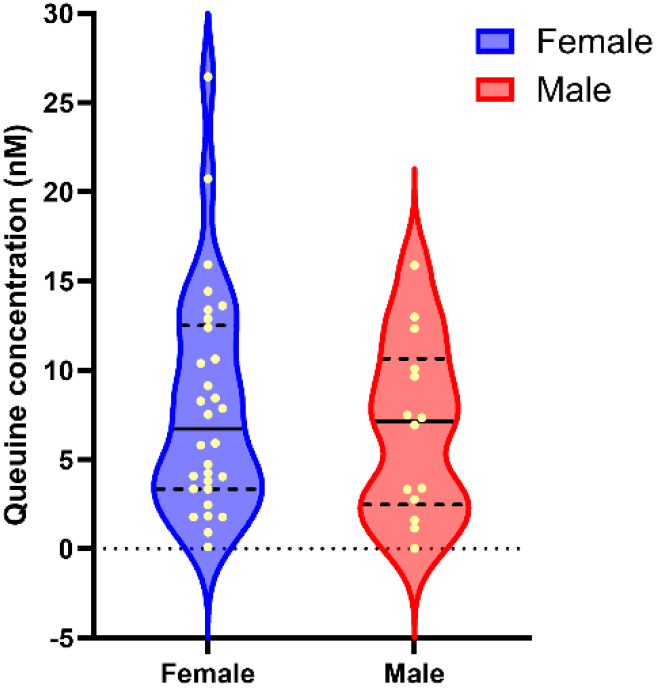
Plasma q levels in healthy male and female subjects

#### d) Carry-over

No deviation from noise was observed following a highest QC injection. Carryover was thus considered negligible in the optimised method. QC levels during the method validation were not affected by injection order.

### Application of the method to clinical human plasma samples

No significant difference in q levels between female (mean=8.0 nM, min = 0.10 nM, max= 9.17 nM and IQR = 8.18 nM) and male (mean = 6.8 nM, min = 0.01 nM, max= 15.89 nM and IQR = 15.88 nM) participants was detected (p = 0.50). Q was not detected in human plasma.

## Discussion

LC-MS/MS-based methods have previously been used to quantify Q-tRNA modification (19). LC-MS/MS has also be developed for measuring as many as 65 other free nucleosides and nucleotides (27), but unfortunately not yet for q nor Q. To our knowledge, there is one study that has analysed q in human plasma with HPLC-MS/MS (15). However, this report did not describe any validation for the method, something that is considered essential for reliable and accurate quantification of clinical samples. Thus, given this limitation and q/Q’s emerging importance in nutrition, physiology and pathology, we present an optimised and validated method which has considered various blood processing approaches, analyte extraction efficiency and quantification for q and Q. This study found that the addition of a stabilization cocktail does not improve the detected levels of q in either plasma or serum, and therefore stabilization cocktail is not necessarily required for sample collection. Either serum or EDTA plasma samples (collected without stabilization cocktail) could be used for measurement of free q in human blood. Serum however, would be preferential because of its lower variability. For extraction, the use of a PBA cartridge is a necessity because it is capable of extracting q at recovery rates greater than 80% in biological samples. The PBA sorbent retains the diol-containing analytes via reversible covalent bonds (28), which can ensure that the nucleic acid successfully binds to the PBA SPE cartridge under alkaline conditions (ammonia solution, pH 11.35) and subsequently this can be eluted by an acidic mobile phase (acetonitrile-water (3:7,v/v) with 5% methanol and 1% formic acid). It should be noted that the extraction recovery for q and Q using the 96-well PBA plate is 10% and 30% lower, respectively, than that using individual PBA cartridges. This is likely due to the smaller amount of sorbent, but clearly there are throughput vs recovery trade-offs to be considered.

The described LC-MS/MS method employed here has been successfully applied by our research group to quantify preQ1, Q-5’MP, Q-3’MP, q and Q in the intracellular extracts and culture medium (9). However, human blood is a more complex matrix, containing high levels of phospholipids, with implications for ion suppression or enhancement in LC-MS/MS analyses (29). Furthermore, q is an endogenous and ubiquitous analyte, for which, and the lack of analyte-free (blank) matrix is a challenge for quantification. The approach using PBS with an additive of 4% BSA to mimic the blank matrix is commonly used to address this challenge (30). However, the surrogate matrix has a potential limitation in that does not contain any anticoagulant (as would be present in actual samples), which could potentially alter the responses of analytes. Another potential limitation is the lack of commercially available isotope-labelled q or Q which could be applied as an internal standard to compensate for the matrix effect, since the matrix effect should affect the relative ionization efficiency similarly for both the analyte and its isotope-labelled internal standard. The synthesis of q and Q is reportedly challenging but the development of commercial isotope-labelled internal standards for these micronutrients would be recommended to progress this field, particularly given the recent guidance of the FDA on this matter (31). In such circumstances the surrogate matrix and biological matrix could be compared. Nonetheless, the validation results here demonstrate this method is accurate and reliable using a surrogate matrix without isotope-labelled standards. The accuracy for Q exceeds the FDA acceptance criteria which should be within ±20% for LLOQ.

To demonstrate the application of this validated method samples from 44 healthy participants were analysed. It was found that q levels in plasma tended to be higher in female subjects than male, but the difference was not statistically significant. The one previously published study found mean plasma q levels to be significantly higher in female volunteers than in males (15). A key observation of the present study was that Q was not detectable in the human plasma samples. It could be the case that q is the major circulating isoform of this micronutrient. It is also possible that all Q metabolites present in food are digested and absorbed as q, and/or that these are processed by the liver, which might only facilitate the entry of q into the blood circulation. It could also be the case that both q and Q enter the blood circulation, but that cells and organs have a stronger absorbative preference for Q. Rapid Q absorption could make it less detectable in blood. Further studies are required to understand the metabolism and pharmacokinetics of q/Q and the method outlined here will be a key enabler for this. Furthermore, with this validated method, q/Q levels in blood samples it is now possible to evaluate the various influences of diet, microbiome and disease. An obviously next step will be to assess circulating levels of the micronutrient in cases of neurodegenerative disease, such as AD, and to assess whether they differ from healthy control subjects.

In conclusion, this method has fulfilled the requirements for a fully validated quantification method for q and Q in biological samples using a surrogate matrix. It was sufficiently validated in terms of accuracy, precision, linearity, recovery, matrix effect and parallelism following the recent FDA guidance. This assay was successfully applied to human plasma samples to support further studies for understanding the physiological and pathophysiological roles of q and Q.

## Ethical approval and consent to participate

The original study protocol was approved by the local ethics committees at the following German institutions: University of Bonn; University of Hamburg; University of Duesseldorf; University of Heidelberg/Mannheim; University of Leipzig and the Technical University of Munich. Written informed consent was obtained from all participants.

## Author contribution statement

XP, SC, BG and JW conceived and designed the studies. XP and SC performed laboratory studies collected and analysed the data. The initial manuscript draft was written by XP. SR, MS, WM, and AR obtained, collected and supplied clinical samples. BG and JW obtained research funding, and managed and supervised the project. All authors commented on manuscript drafts and approved the final manuscript.

