## Supplemental data for "Development, validation and application of an LC-MS/MS method quantifying free forms of the micronutrients queuine and queuosine in human plasma using a surrogate matrix approach"

**Supplementary method**

The plasma from these 44 persons were selected from the German study on Aging, Cognition and Dementia (AgeCoDe) biobank (Ramirez et al., 2015, doi:10.3233/JAD-141521). The original study protocol was approved by the local ethics committees at the following German institutions: University of Bonn; University of Hamburg; University of Duesseldorf; University of Heidelberg/Mannheim; University of Leipzig and the Technical University of Munich. Written informed consent was obtained from all participants. The main assessment instrument at all visits included the Structured Interview for Diagnosis of Dementia of Alzheimer type, Multi-infarct Dementia and Dementia of other etiology according to DSM-IV and ICD-10 (SIDAM), and diagnosis of AD was established according to the NINCDSADRDA criteria for probable AD (McKhann et al., 1984, doi:10.1212/wnl.34.7.939; Zaudig et al., 1991, doi:10.1017/s0033291700014811).

AgeCoDe is a longitudinal study, where participants were recruited in primary care centers in six German cities. Inclusion criteria were to be at least 75 years old and cognitively healthy according to the general practitioner’s judgment. Every ~18 months interval participants are followed up with personal interviews and neuropsychological assessments. Nine follow-ups (FUs) were completed. Blood samples for EDTA plasma were obtained at the third visit, processed, and stored at -80°C. For this study, the third visit is considered the baseline. Healthy participants from AgeCoDe included in this study remained cognitively unimpaired until the last FU.

|  | **MCX 1cc** | **WCX 1cc** | **SAX 3cc** | **HLB 1cc** | **ENVI CARB 3cc** | **PBA 1cc** |
| --- | --- | --- | --- | --- | --- | --- |
| **Condition** | Methanol | Methanol | Methanol | Methanol | Methanol | 1% FA in  ACN: Water  (1:1) |
| **Equilibrate** | Water | Water | Water | Water | Water | 1mL ammonia solution pH 11.35 |
| **Load** | Spiked sample with 0.8% acetic acid | Spiked sample with 0.8% acetic acid | Spiked sample diluted with PBS pH 7 | Spiked sample diluted with 4% H_3_PO_4_ in H2O (1:1)  pH 4.92 | Spiked sample diluted with PBS pH 7 | Spiked sample diluted with ammonia solution pH 11.35 |
| **Wash** | 0.1M HCl | 0.1M HCl | 50mM sodium acetate in 5% methanol pH 7 | 5% methanol in water | Water | 5% ACN with ammonia solution pH 11.35 |
| **Elute** | 5% NH_4_OH in methanol | 5% NH_4_OH in methanol | 2% formic acid in methanol | 100% methanol | Methanol +  dichloromethane  +ethyl acetate | 1% FA in ACN: water (3:7) + 5% methanol |
| **Recovery** | Q – 54.0%  q – 44.5% | Q – 0.4%  q – 0.2% | Q – 3.8%  q – 4.2% | Q – 6.0%  q – 1.3% | Q – nil  q – nil | Q – 71.0%  q – 82.0% |

**Supplementary Table 1** SPE phases tested and protocols for extraction q and Q
